# Relating pupil diameter and blinking to cortical activity and hemodynamics across arousal states

**DOI:** 10.1101/2022.06.15.496309

**Authors:** Kevin L. Turner, Kyle W. Gheres, Patrick J. Drew

## Abstract

Arousal state affects neural activity and vascular dynamics in the cortex, with sleep associated with large changes in the local field potential (LFP) and increases in cortical blood flow. We investigated the relationship between pupil diameter and blink rate with neural activity and blood volume in the somatosensory cortex in male and female unanesthetized, head-fixed mice. We monitored these variables while the mice were awake, during periods of rapid eye movement (REM), and non-rapid eye movement (NREM) sleep. Pupil diameter was smaller during sleep than in the awake state. Changes in pupil diameter were coherent with both gamma-band power and blood volume in the somatosensory cortex, but the strength and sign of this relationship varied with arousal state. We observed a strong negative correlation between pupil diameter and both gamma-band power and blood volume during periods of awake rest and NREM sleep, though the correlations between pupil diameter and these signals became positive during periods of alertness, active whisking, and REM. Blinking was associated with increases in arousal and decreases in blood volume when the mouse was asleep. Bilateral coherence in gamma-band power and in blood volume dropped following awake blinking, indicating a ‘reset’ of neural and vascular activity. Using only eye metrics (pupil diameter and eye motion), we could determine the mouse’s arousal state (‘Awake’, ‘NREM’, ‘REM’) with greater than 90% accuracy with a 5 second resolution. There is a strong relationship between pupil diameter and hemodynamics signals in mice, reflecting the pronounced effects of arousal on cerebrovascular dynamics.

**Significance Statement:** Determining arousal state is a critical component of any neuroscience experiment. Pupil diameter and blinking are influenced by arousal state, as are hemodynamics signals in the cortex. We investigated the relationship between cortical hemodynamics and pupil diameter and found that pupil diameter was strongly related to the blood volume in the cortex. Mice were more likely to be awake after blinking than before, and blinking ‘resets’ neural activity. Pupil diameter and eye motion can be used as a reliable, non-invasive indicator of arousal state. As mice transition from wake to sleep and back again over a timescale of seconds, monitoring pupil diameter and eye motion permits the non-invasive detection of sleep events during behavioral or resting-state experiments.

## Introduction

The dynamics of the eyes convey information about mental state. Besides their respective roles in controlling light levels and protecting the eye, pupil diameter and blinking give information about the state of neural activity in the brain (Strauch et al., 2022). However, for measures of pupil diameter and blinking to be useful, we must understand their relationship to neural and vascular physiology. Pupil dilations are associated with higher levels of arousal during the awake state (Drew et al., 2001; Hess and Polt, 1964; Kahneman and Beatty, 1966; Morad et al., 2000; Onorati et al., 2013; Yoss et al., 1970) and correlate with increased sympathetic activity (Bradley et al., 2008). The pupil will usually dilate when a subject is performing a task or making a decision (Burlingham et al., 2022; Einhauser et al., 2010; Gilzenrat et al., 2010; Hakerem and Sutton, 1966; Nassar et al., 2012) and recent rodent work has shown that pupil diameter fluctuations temporally track cortical state (McGinley et al., 2015; Reimer et al., 2014; Vinck et al., 2015). In both rodents and primates, pupil dilation reflects noradrenergic tone in the cortex as well as activity in the locus coeruleus (LC) (Joshi et al., 2016; Larsen and Waters, 2018; Preuschoff et al., 2011; Reimer et al., 2016). In addition to its effects on neurons, noradrenergic input from the LC causes vasoconstriction of cortical arteries (Bekar et al., 2012). LC activity and noradrenergic tone provides a potential mechanism for coupling arousal to cortical hemodynamics (Pisauro et al., 2016), as changes in pupil diameter are also correlated with blood-oxygen-level-dependent (BOLD) signals seen in neuromodulatory centers during functional magnetic resonance imaging (fMRI) (Pais-Roldan et al., 2020; Sobczak et al., 2021). Both head-fixed and freely behaving mice frequently sleep with their eyes open (Karimi Abadchi et al., 2020; Senzai and Scanziani, 2022; Turner et al., 2020; Yuzgec et al., 2018). Pupil diameter decreases during sleep and can be used to detect sleep events on a minute-to-minute timescale (Karimi Abadchi *et al.*, 2020; Yuzgec *et al.*, 2018), making it a particularly useful metric for attention and behavioral monitoring.

Blink rate is influenced by mental state and fatigue (Holland and Tarlow, 1972; Nakamori et al., 1997; Stern et al., 1994; Van Orden et al., 2001). Humans blink every few seconds (Stern et al., 1984), though blinking is less frequent in rodents (Kaminer et al., 2011). Blink rate is modulated by dopaminergic tone (Karson, 1983), and is higher in individuals with schizophrenia (Karson, 1979; Karson et al., 1990). Blinking causes brief decreases in neural activity in visual areas that are similar to transient darkening (Gawne and Martin, 2000; Golan et al., 2016). However, blinking also drives BOLD signals in the visual cortex (Bristow et al., 2005a; Bristow et al., 2005b; Hupe et al., 2012) and somatosensory regions (Guipponi et al., 2015), and blinks are correlated with BOLD activity in the default mode network (Nakano et al., 2013). Blinking is correlated with changes in neural and vascular dynamics across the brain, though the correlates of blinking with arousal are not as well understood.

To better understand how pupil diameter and blinking relate to neural activity and cortical hemodynamics across arousal states, we analyzed video of the eye with concurrent monitoring of neural activity and blood volume in the somatosensory cortex of head-fixed mice of both sexes. We investigated the relationship between spontaneous changes in pupil diameter and these physiological signals during the awake state, as well as during REM and NREM sleep. We also tracked how neural activity and cortical blood volume changed around blinking events. Using only changes in pupil diameter and eye position, we found that we can accurately detect and categorize REM and NREM sleep events on a time scale of seconds, providing a simple and robust measure of arousal for studies employing eye monitoring in rodents.

## Materials and Methods

Data presented here is from 22 C57BL/6J mice (12 males, Jackson Laboratory, Bar Harbor, ME) between the ages of 3 and 8 months of age. Data from 14 of these mice was previously published (Turner *et al.*, 2020), and we have added imaging/recordings from an additional 8 mice. We obtained a total of 442.75 hours of data (20.1 ± 5.3 hours per mouse) of naturally occurring ‘Awake’ (61.4 ± 17.3 %), ‘NREM’ sleep (34.2 ± 16.1%), and ‘REM’ sleep (4.4 ± 2.8%) data.

### Arousal state nomenclature

Capitalized italics (*Rest, NREM, REM, Alert, Asleep,* and *All*) denote arousal states with specific inclusion criteria. Capitalized non-italics with single quotes refer to individual 5-second labels of arousal state classified using a machine learning algorithm (‘Awake’, ‘NREM’, ‘REM’) and we use the term ‘Asleep’ (non-italicized) to refer to events classifications as either ‘NREM’ or ‘REM’ (Note **Fig. 4** and **Fig. 5**). When generally discussing arousal state in all other contexts, we use the terms awake, sleep, NREM sleep, REM sleep, etc. with no italics or other indicators.

### Animal procedures

This study was performed in accordance with the recommendations in the Guide for the Care and Use of Laboratory Animals of the National Institutes of Health. All procedures were performed in accordance with protocols approved by the Institutional Animal Care and Use Committee (IACUC) of The Pennsylvania State University (Protocol # 201042827). A head-bar, as well as cortical, hippocampal, and nuchal muscle electrodes along with bilateral polished and reinforced thinned-skull windows (Drew et al., 2010; Shih et al., 2012; Zhang et al., 2022b) were surgically implanted under isoflurane anesthesia (5% induction, 2% maintenance). Detailed surgical procedures have been previously described (Mirg et al., 2022a; Mirg et al., 2022b; Turner *et al.*, 2020; Winder et al., 2017). Following surgery, animals were housed individually on a 12-hr. light/dark cycle (lights on at 7:00 am) with food and water *ad libitum.* Each animal was gradually acclimated to head-fixation in the weeks following recovery. Following the conclusion of imaging experiments, animals were deeply anesthetized and transcardially perfused with heparin-saline followed by 4% paraformaldehyde for histological verification of electrode placement (Adams et al., 2018; Drew and Feldman, 2009).

### Physiological data acquisition

Data were acquired with a custom LabVIEW (National Instruments, Austin, TX) program (https://github.com/DrewLab/LabVIEW-DAQ). For details on intrinsic optical signal (IOS) imaging, electromyography (EMG), electrophysiology, whisker stimulation, and behavioral measurements see (Turner *et al.*, 2020; Zhang *et al.*, 2022b). Previous work from our lab has shown that the intrinsic signal is not affected by skull/brain movement or other motion artifacts. In previously published experiments (Winder *et al.*, 2017), we looked at reflectance changes in a piece of clay mounted over the cranial window, which would be sensitive to any motion artifacts as well as any other non-hemodynamic noise sources. The reflectance changes were on the order of 0.01%, much smaller than the ~20% changes in reflectance (ΔR/R) we see during sleep. Furthermore, 2-photon imaging from our lab of mice running on a treadmill (Gao and Drew, 2016; Echagarruga et al.,. 2020) showed ~2 μm of brain/skull motion during locomotion (a more extreme imaging condition than presented here), which is too small to impact the IOS. IOS reflectance was converted to changes in total hemoglobin (Δ[HbT]) using the Beer-Lambert law (Ma et al., 2016a; Ma et al., 2016b). Data was acquired in 15-minute intervals with a < 1-minute gap in between for saving data to disk. The vibrissae (left, right, or a third air puffer not directed at the body as an auditory control) were randomly stimulated with air puffs (0.1 seconds, 10 pounds-force per square inch (PSI)) occurring every 30–45 second for the first ~1 hr. of imaging.

### Pupil diameter measurement

The pupil was illuminated with 780 nm light (measured at 0.02–0.05 mW/mm^2^, Thorlabs S120VC) as mice are functionally blind to these wavelengths (Breuninger et al., 2011; Chang et al., 2013). To prevent constriction of the pupil by the illumination for IOS, a dichroic mirror located above the head was used to block as much of the IOS illumination from the eyes as possible so that the mouse was only exposed to a faint glow in its periphery (0.002–0.005 mW/mm^2^) with no visible wavelength light shining directly into its eyes. We did not try to eliminate all green light reaching the eye because green light has sleep-promoting effects (Pilorz et al., 2016). While we did not quantify the pupil diameter in complete darkness as higher baseline dilations would result in the pupil edges being obscured by the eyelid. All pupil diameter measurements were verified by manual inspection (KLT).

### Data Availability

Data and sample files for running the pupil tracking algorithm are available at https://datadryad.org/stash/share/pv4ZmJnSk65Y6yWxoO6jdb9ou5H-x5wfOJOTCPkBntE and analysis code is available at https://github.com/KL-Turner/Turner-Manuscript2022. Data was analyzed with code written by K.L.T, K.W.G, and P.J.D (MathWorks, MATLAB 2019b-2022a, Natick, MA).

### Automated pupil diameter measurement and blink detection

Pupil diameter was extracted from videos of the eye taken using a Basler GigE camera (acA640-120gm, Ahrensburg, Germany) with a 75 mm double Gauss, fixed focal length lens (#54-691, Edmond Optics Barrington, NJ) at 30 frames/second. Our pupil detection algorithm was adapted from the thresholding in Radon space (TiRS) algorithm developed to determine vessel cross-sections (Gao and Drew, 2014). The sclera and pupil were defined as the area within a user-selected region of interest (ROI) created by outlining the first frame with the eye fully open. Images were inverted so that the pupil had the maximal intensity and were 2-D median filtered ([5,5] x,y pixel median). The pixel intensity distribution was fit with a normal distribution using maximum likelihood estimation (MLE), and pixels above a user-defined threshold (average of 1 ± 0.25 standard deviations from the MLE mean) were set to 1 and all other pixel’s intensities were set to 0. The binarized image was then converted to Radon space and normalized within each projection angle (Gao and Drew, 2014). A second threshold was applied prior to conversion back to image space and any holes within the filtered pupil object were filled. Frame-wise pupil area was then calculated using a boundary classification algorithm. Occasional obstructions of the pupil (by a vibrissae) were identified by detecting rapid fluctuations in pupil area. For these frames, the threshold used for thresholding in Radon space was iteratively decreased until the pixel area was within frame-wise change boundaries. Any periods where the pupil was obscured or otherwise unmeasurable were discarded from analysis. The TiRS algorithm was developed to detect small changes in the area of an ellipse (Gao and Drew, 2014). Other techniques, including using a deep neural network (DNN) such as DeepLabCut (Mathis et al., 2018), can be used to track the pupil diameter and position as well (Privitera et al., 2020), though this requires training the network. While the pupil was slightly elliptical in appearance due to the angle of the camera and movement of the eye, the correlation between the major and minor axis of the object’s area was 0.96 ± 0.02 (N = 22 mice), indicating the ellipse at the pupil boundary does not appreciably change shape, only size. The area (A) was used to calculate the diameter (d) of the pupil using the formula 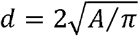. MATLAB function(s): roipoly, medfilt2, imcomplement, mle, pdf, radon, iradon, bwboundaries, bwconvhull, imfill, regionprops, diff, fillmissing.

### Blink detection

Blinking was detected independently of pupil diameter from the same ROI as used for pupil diameter measurements. Rapid changes in eyelid motion were detected by using the sum of pixel intensities within the user-defined ROI. The intensity of all pixels for each frame (sclera and pupil) were summed, and a frame-to-frame ROI intensity difference was calculated. As the eyelid obscures the pupil during blinks, rapid changes in overall pixel intensity corresponded to eyelid movements. A threshold was applied to binarize changes in luminance across frames, and blinking events were extracted as above thresholded changes in image intensity. As mice frequently blinked several times in rapid succession, blinking events that occurred with less than 1 second between them were concatenated into a single blinking bout. Periods where a false-positive blink was detected, such as from the eye only partially closing, were also discarded. All detected blinking events were verified by manual inspection (KLT).

### Electrophysiological analysis

Discrete LFP bands were digitally bandpass filtered from the broadband data using a third-order Butterworth filter into the following bands: delta [1–4 Hz], theta [4–10 Hz], alpha [10–13 Hz], beta [13–30 Hz], gamma [30–100 Hz]. The filtered signal was then squared, low-pass filtered < 10 Hz, and resampled at 30 Hz. Time frequency spectrograms were calculated using the Chronux toolbox (Bokil et al., 2010), version 2.12 v03, function mtspecgramc with a 5 second window and 1/5 second step size using [5,9] tapers and a passband of 1-100 Hz to encompass the LFP. Electromyography (EMG, 300 Hz – 3 kHz) from the nuchal (neck) muscles was bandpass filtered, squared, convolved with a Gaussian kernel with 0.5 second standard deviation, log transformed, and then resampled at 30 Hz. MATLAB function(s) butter, zp2sos, filtfilt, gausswin, log10, conv, resample.

### Sleep scoring

Sleep states were scored consistent with previously published criteria (Cirelli, 2009; Saper and Fuller, 2017; Weber and Dan, 2016). NREM sleep is marked by predominantly elevated cortical delta-band power and lower EMG power during slow-wave (NREM sleep) (Amzica and Steriade, 1998; Steriade et al., 1993). REM sleep is marked by elevated hippocampal theta-band power and elevated cortical gamma-band power with even further reduced EMG power (muscle atonia) (Cantero et al., 2004; Le Van Quyen et al., 2010; Montgomery et al., 2008; Sullivan et al., 2014). Periods of user-verified awake rest greater than 5 second in duration with no whisker stimulation, no whisker motion, and no detectable body motion were identified and used baseline characterization of all signals as well as for z-scoring the pupil. Sleep scoring was performed as in (Turner *et al.*, 2020). Every 5 second interval was classified as either Awake, NREM sleep, or REM sleep using a bootstrap aggregating random forest model with the predictors of cortical delta LFP, cortical beta LFP, cortical gamma LFP, hippocampal theta LFP, EMG power, heart rate, and whisking duration. Sleep model accuracy was validated using the out-of-bag error during model training. MATLAB function(s): TreeBagger, oobError, predict.

### Pupil diameter during different arousal states

Pupil diameter was taken from awake resting events (*Rest*) (≥ 10 seconds in duration), volitional whisking (*Whisk*) (2–5 seconds), whisker stimulation (*Stim*) (0.1 seconds, 10 PSI to vibrissa), *NREM* (≥ 30 seconds), and *REM* (≥ 60 seconds). Pupil diameter was low-pass filtered <1 Hz with a fourth-order Butterworth filter. Changes in whisking-evoked and stimulus-evoked (contralateral, auditory) diameters were taken as the change in diameter relative to the mean of the 2 seconds preceding the event onset. Classifications of *Alert* or *Asleep* were taken as 15-minute periods with no whisker stimulation and at least 80% of a given classification (Awake for *Alert,* NREM or REM for *Asleep*) within a 15-minute recording. The classification of *All* denotes all data taken during periods with no sensory stimulation, independent of arousal state. MATLAB function(s): butter, zp2sos, filtfilt.

### Power spectra and coherence

Spectral power was estimated using the Chronux toolbox (Bokil *et al.*, 2010) function mtspectrumc. For pre-whitening spectra, the first derivative was taken of the mean-subtracted data before the power calculation. Gamma-band power measurements were scaled by a factor of 1×10^19^ so that the magnitude of the neural changes (arbitrary units, a.u.) were more in line with those from the hemodynamic signal for ease of comparison. Coherence analysis was run for each data type using the Chronux function coherencyc. MATLAB function(s): detrend, diff.

### Δ[HbT]/Gamma-band power vs. pupil relationship

Mean changes in total hemoglobin (Δ[HbT]) or gamma-band power (ΔP/P), and pupil diameter (z-units) during each arousal state classification (‘Awake’, ‘NREM’, ‘REM’) was plotted as a 2D histogram to highlight clustering in each class. Each classification was assigned a color and the three images were merged as a composite in Fiji (ImageJ). MATLAB function(s): histogram2.

### Cross-correlation

Data was low-pass filtered < 1 Hz using a fourth-order Butterworth filter and mean-subtracted. Cross-correlation analysis was run for each arousal state with either a ± 5 second lag (*Rest, NREM, REM*) or ± 30 second lag (*Alert, Asleep, All*) depending on duration of the behavioral state. MATLAB function(s): xcorr, filtfilt, detrend.

### Interblink interval and blink-associated physiology

Due to gaps in recording to save the data to disc, interblink interval was calculated between blinks occurring within 15-minute records and not blinks on the edges of trials. Blinks that occurred with 1 second of each other were linked together as blinking bouts, and all blink-triggered analysis were with respect to the first blink in a series. Blink-triggered averages were separated into two groups depending on the arousal state classification of the 5 second bin prior to the blink, that being either Awake (arousal state classification of Awake) or Asleep (arousal state classification being either NREM or REM).

### Probability of sleep as a function of pupil diameter

The probability of being in each arousal state (‘Awake’, ‘NREM,’ ‘REM’) as a function of pupil diameter was obtained from the mean diameter during each 5 second sleep score and binning the value into a histogram, which was then smoothed with a median filter and Savitzky–Golay filter. MATLAB function(s): medfilt1, sgolayfilt.

### Tracking of pupil location

Motion of the pupil was tracked by using the X and Y coordinates of the centroid obtained during pupil tracking and comparing the change in position to the baseline position during *Rest*. Frame-by-frame changes in centroid location were low-pass filtered < 10 Hz using a 4^th^-order Butterworth filter. Motion of the eye was evaluated during each arousal state classification (Awake, NREM, REM) by taking the cumulative sum of the absolute change in centroid position within each 5 second bin. Transitions between arousal states (‘Awake’, ‘NREM’, ‘REM’) for pupil diameter, position, and motion were extracted from arousal classifications with 30 seconds of consecutive classifications of one state followed by 30 seconds of another.

### Eye, physiological, and combined model comparison

The Eye model was trained using only eye metrics, pupil diameter (mean, variance, minimum of both z-unit and mm diameter), position (mean, variance, and maximum displacement of the x,y centroid), and motion (sum, variance of centroid’s absolute velocity) from data with open eyes. The Physiological model was trained using non-eye-based measures (cortical delta power, hippocampal theta power, EMG, etc. - see (Turner *et al.*, 2020)) from the same data so that the training/testing sets were identical among models. The Combined model used a union of all parameters from both models. Each model used was trained using a bagged random forest composed of 128 decision trees, with out-of-bag error and confusion matrices from each animal being calculated at the time of model training using a 70-30 (training-testing) split of randomly shuffled labels taken from each arousal state. MATLAB function(s): TreeBagger, oobError, predict, confusionchart.

### Experimental design and statistical analysis

Researchers were not blind to the experimental conditions or data analysis. Sample sizes are consistent with those of previously published studies (Echagarruga et al., 2020; Huo et al., 2015; Turner *et al.*, 2020; Winder *et al.*, 2017). Statistical evaluations were made using either generalized linear mixed-effects (GLME) models with the arousal state as a fixed effect, mouse identity as a random effect, and hemisphere (L/R, if applicable) as an interaction with the animal ID; or using a paired t-test where appropriate. Unless otherwise stated, statistical results report p-values from a GLME test. All reported quantifications are mean ± standard deviation unless otherwise indicated. Unless otherwise noted, all pupil diameter measurements are in z-units. MATLAB function(s): fitglme, ttest.

## Results

Unanesthetized mice were head-fixed under an IOS imaging setup (Huo et al., 2014; Pisauro et al., 2013; Sirotin and Das, 2009; Vazquez et al., 2014; Winder *et al.*, 2017) with concurrent electrophysiology to measure changes in neural activity from the vibrissa region of the somatosensory cortex (Petersen, 2007; 2014) and hippocampal CA1. We also tracked whisker motion (Winder *et al.*, 2017), electromyography of the nuchal muscles (Turner *et al.*, 2020), and pupil diameter/blinking (Larsen and Waters, 2018; Reimer *et al.*, 2014; Reimer *et al.*, 2016; Vinck *et al.*, 2015) **(Fig. 1a**). Using the thresholding in Radon space algorithm (Gao and Drew, 2014), we detected the outline of pupil from video frames (**Fig. 1b-e, Video 1**) in order to quantify diameter changes. Blinks were detected from rapid pupil diameter changes (**Fig. 1f**). During the awake state, there are frequent bouts of whisker motion that are correlated with increases in pupil diameter, and the EMG has high power. The LFP has reduced low-frequency power and hemodynamic fluctuations have low amplitude, except during periods of extended behavior or sensory stimulation. During NREM sleep, whisker motion and EMG power are much lower and the pupil diameter is smaller than in the awake state. Cortical LFP and hemodynamic signals both begin to increase in amplitude with low frequency oscillations in broadband LFP power. During REM sleep, whisker motion increases along with eye movement, but the pupil remains constricted. Power in the EMG is at its lowest point due to muscle atonia (with occasional twitches), blood volume increases substantially above both awake and NREM levels (Bergel et al., 2018), and a prominent theta-band is visible in the hippocampal LFP. A representative example of how pupil diameter changes along with fluctuations in Δ[HbT], hippocampal LFP, and other behavioral cues during each arousal state is shown in **Fig. 1f** and **Video 1.** Since mice have been shown to sleep with their eyes open during both head-fixed (Karimi Abadchi *et al.*, 2020; Turner *et al.*, 2020; Yuzgec *et al.*, 2018) and freely moving (Senzai and Scanziani, 2022) conditions, we tested whether neural activity and hemodynamics might differ between the eyes-open and eyes-closed instances of the REM state. We compared the eyes-open REM sleep physiology with that during obtained during a few instances of REM sleep with eyes-closed in a subset of mice (N = 8) that was excluded from subsequent analyses due to the inability to measure pupil diameter. The change in total hemoglobin (Δ[HbT]) during REM with eyes open (73.9 ± 12.9 μM) was not significantly different from the hemoglobin changes during REM sleep with eyes closed (76.2 ± 21.2 μM, *p* < 0.49, paired *t*-test). The difference in theta-band power in the hippocampus during eyes-open REM and eyes-closed REM was also not statistically significant, (1.06 ± 1.11 A.U. vs. 1.06 ± 1.10 A.U., respectively, *p* < 0.96, paired *t*-test). The lack of detectable difference between the eyes-open and eyes-closed sleep state suggests that whether or not they eyes are open has little impact on sleep physiology.

**Fig. 1.**
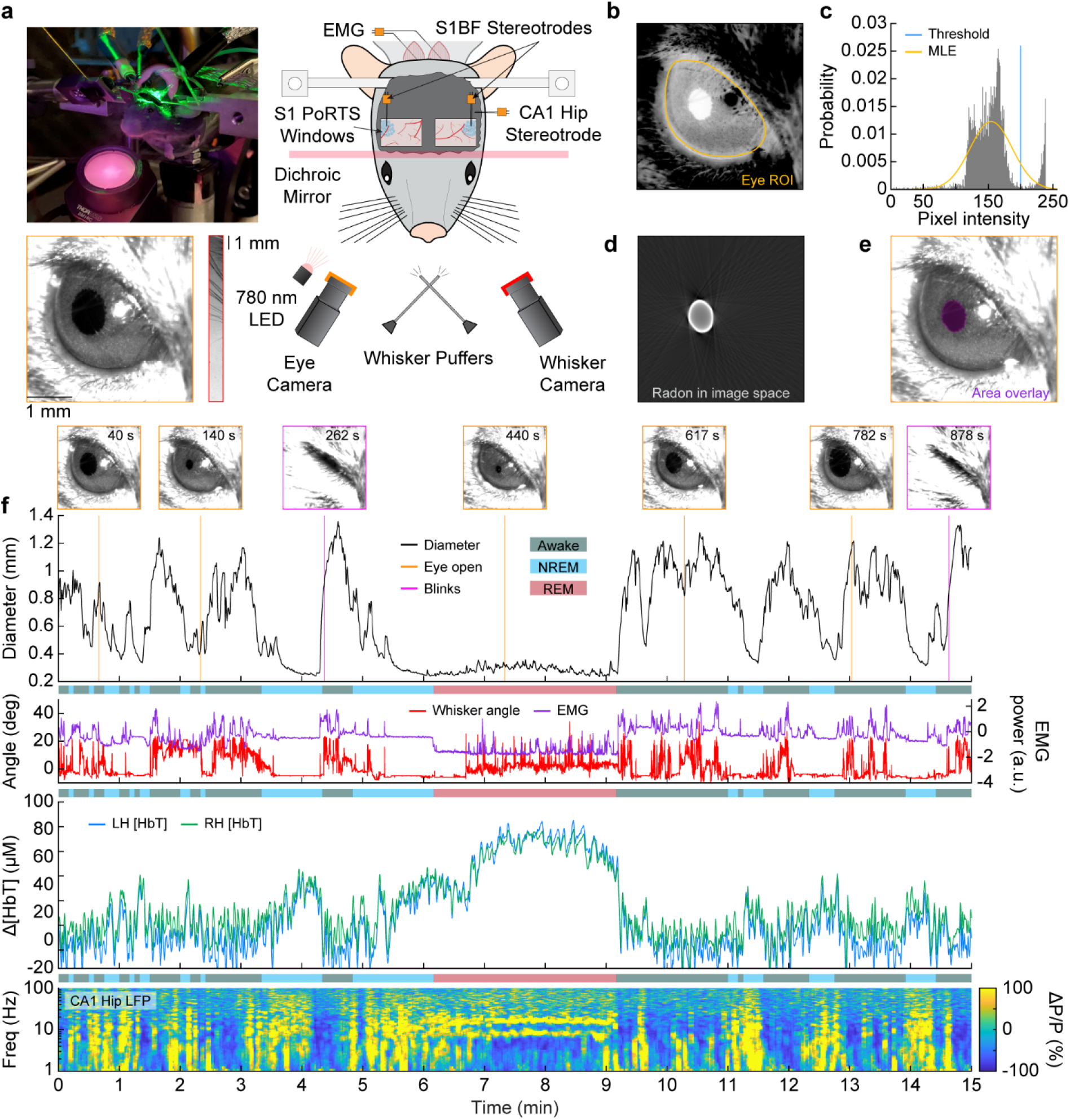
Pupil diameter tracks arousal state. **(a)** Photograph and schematic of the experimental setup. IOS imaging of the whisker portion of somatosensory cortex collected using illumination at an isosbestic point of hemoglobin (530 nm or 568 nm) through bilateral thinned-skull windows. Electrodes were implanted into layer 5 of the somatosensory cortex and into hippocampal CA1. EMG electrodes were also implanted into the nuchal muscles of the neck. A camera is directed towards the whiskers to track their movement. Another camera captures the eye, which is illuminated by infrared 780 nm light, to observe pupil diameter and blinking. Directed air puffers allow controlled stimulation of the left and right whiskers. **(b)** ROI around the mouse’s eye. **(c)** Histogram of pixel intensities in the ROI showing separation between the intensities in the pupil and sclera. A threshold (blue) is set for initial diameter estimation based on a maximum likelihood estimate of the sclera intensity (MLE, orange). **(d)** Image of the pupil after has been thresholded in Radon space and transformed back into image space. **(e)** Detected pupil superimposed on video frame. **(f)** Example showing changes in pupil diameter, whisker motion, EMG power, Δ[HbT], and CA1 LFP during awake, NREM, and REM periods.

### Quantification of pupil diameter across arousal states

To quantify how fluctuations in pupil diameter change with arousal state, we compared the pupil’s diameter during several distinct arousal states and behaviors. Pupil diameter was largest during periods of arousal and smallest during sleep. To standardize measures across animals and to account for slight differences in baseline illumination intensity, we z-scored the pupil diameter to the mean and standard deviation during all periods of awake quiescence lasting at least 5 seconds in length. Italics denote periods meeting our arousal state criteria. 15-minute periods with at least 80% of its model scores as ‘Awake’ were classified as *Alert.* 15-minute periods with 80% of the time in ‘NREM’ or ‘REM’ sleep were classified as *Asleep. All* was all data, regardless of arousal state. Periods with whisker stimulation were excluded from all states. Two mice did not have any 15-minute periods that met the criteria for the *Alert* or *Asleep* categories (N = 20 mice included) with the other four behaviors (*Rest, NREM, REM, All*) all having N = 22 mice. *Rest,* defined as all periods of awake quiescence lasting at least 10 seconds in length, had a mean pupil diameter of 0.6 ± 0.17 mm **(Fig. 2a)** or −0.25 ± 0.88 z-units **(Fig. 2b)**. We used a GLME model to compare the pupil’s diameter among states, using the Bonferroni correction for multiple comparisons (10), which puts the adjusted significance threshold (α) at 0.005. The mean z-unit during *Rest* was non-zero because the minimum duration used for z-scoring (5 seconds) was shorter than the minimum duration for inclusion in *Rest* (10 seconds). During volitional whisking bouts lasting 2–5 seconds, pupil diameter increased to 0.89 ± 0.17 mm (*p* < 2.3×10^-19^ versus *Rest*) or 3.39 ± 0.88 z-units (*p* < 2.1×10^-22^). Following brief stimulation of the vibrissae, pupil diameter increased to 0.75 ± 0.18 mm (*p* < 9.2×10^-8^) or 1.95 ± 1.82 z-units (*p* < 1.9×10^-11^). The pupil diameter decreased during *NREM* (> 30 seconds in length) to 0.35 ± 0.07 mm (*p* < 8.1×10^-16^) or −3.43 ± 0.64 z-units (*p* < 6.2×10^-19^) and even further during *REM*(≥ 60 seconds in length) to 0.26 ± 0.03 mm (*p* < 3.2×10^-23^) or −4.56 ± 0.75 z-units (*p* < 2.3×10^-27^). Additional statistical comparisons for the difference in pupil size between each arousal state can be found in **Table 1** and **Table 2**. Periods of volitional whisking and whisker stimulation caused increases in pupil diameter of 1.44 ± 0.47 z-units and 1.61 ± 0.62 z-units, respectively. Auditory controls dilated the pupil by 0.98 ± 0.58 z-units **(Fig. 2c)**. Note that while the z-unit diameter was set relative to awake resting events, these increases are with respect to changes in z-unit diameter relative to the 2 seconds prior to event onset, which was a mixture of awake resting and volitional behaving data of various durations.

**Fig. 2.**
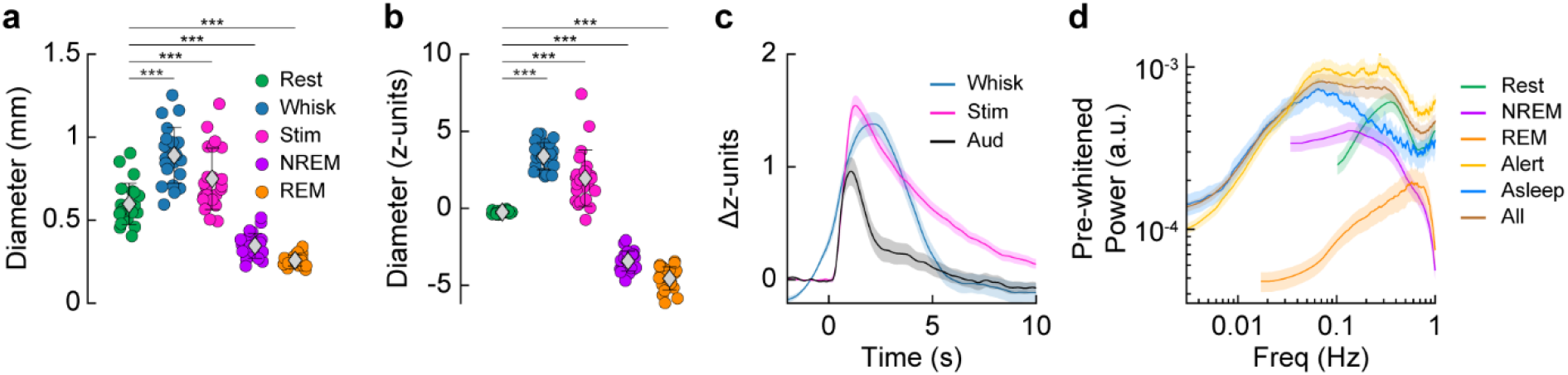
Quantification of pupil diameter across arousal states. **(a)** Mean pupil diameter (mm) for each arousal state. Mean of each animal is a circle. Grey diamond and error bars are mean ± std. across animals for each arousal state. **(b)** Mean pupil diameter (z-units) for each arousal state. **(c)** Whisking-evoked and stimulus evoked increases in pupil diameter based on the 2 seconds prior to event onset. **(d)** Pre-whitened power spectrum of pupil diameter (z-units) during different arousal states. Shading **(c, d)** denotes standard error of the mean. Error bars **(a, b)** denote standard deviation. Statistical comparisons shown are between *Rest* and other states using a GLME model with Bonferroni correction for multiple comparisons (10) * α < 0.005, **α < 0.001, ***α < 0.0001. See **Table 1** and **Table 2** for additional statistical comparisons between each arousal state in **(a, b)**.

**Table 1.** Statistical comparison of Mean pupil diameter across arousal states (mm)

|  | Whisk | Stim | NREM | REM |
| --- | --- | --- | --- | --- |
| Rest | ***$2.3 \times 10^{-19}$ | ***$9.2 \times 10^{-8}$ | ***$8.1 \times 10^{-16}$ | ***$3.2 \times 10^{-23}$ |
| Whisk | | ***$5.4 \times 10^{-7}$ | ***$1.2 \times 10^{-38}$ | ***$2.8 \times 10^{-44}$ |
| Stim | | | ***$2.3 \times 10^{-28}$ | ***$6.0 \times 10^{-35}$ |
| NREM |  |  |  | *0.0012 |

Bonferroni corrected significance levels (10): \* $\alpha < 0.005$ , \*\* $\alpha < 0.001$ , \*\*\* $\alpha < 0.0001$

**Table 2.** Statistical comparison of Mean pupil diameter across arousal states (z-units)

|  | Whisk | Stim | NREM | REM |
| --- | --- | --- | --- | --- |
| Rest | ***$2.1 \times 10^{-22}$ | ***$1.9 \times 10^{-11}$ | ***$6.2 \times 10^{-19}$ | ***$2.3 \times 10^{-27}$ |
| Whisk | | ***$3.1 \times 10^{-6}$ | ***$2.3 \times 10^{-43}$ | ***$2.2 \times 10^{-49}$ |
| Stim | | | ***$1.2 \times 10^{-34}$ | ***$1.3 \times 10^{-41}$ |
| NREM |  |  |  | **0.00019 |

Bonferroni corrected significance levels (10): \* $\alpha < 0.005$ , \*\* $\alpha < 0.001$ , \*\*\* $\alpha < 0.0001$

To look at changes in the power spectrum of the pupil diameter during different arousal states, we pre-whitened the pupil diameter throughout analysis during periods of continuous *Rest* (> 10 second duration), continuous *NREM* (> 30 second duration), continuous *REM* (> 60 second duration), and 3 additional longer-length classifications (*Alert, Asleep,* and *All*) by taking the first temporal derivative of the z-unit diameter prior to power estimation. Except for *REM,* the pre-whitened power spectra of the pupil diameter during the longer length behaviors were similar to each other at the lower frequencies **(Fig. 2d)**. These findings support previous reports that the pupil diameter increases with arousal and attention but decreases during periods of sleep (Karimi Abadchi *et al.*, 2020; Yuzgec *et al.*, 2018).

### Relationship between pupil diameter and hemodynamic and neural signals

We next asked how well the pupil diameter correlated with both hemodynamic and neural signals in the somatosensory cortex of mice both during awake behaviors and during different sleep states. For each mouse, the pupil diameter was compared to both hemispheres’ neural and hemodynamic signals, yielding 2*N measurements for each behavior. The non-independence (due to within animal correlations) was accounted for as an interaction term in the statistical comparison (see Materials and Methods). During the ‘Awake’ state, changes in pupil diameter and gamma-band power were small. When the animals were in ‘REM’ and ‘NREM’ sleep, the pupil constricted and gamma-band power increased substantially relative to the ‘Awake’ state (**Fig. 3a**). We then looked at the coherence and the cross-correlation between the envelope of gamma-band power (ΔP/P) and pupil diameter. The minimum cross-correlation during *Rest* was −0.16 ± 0.20 at a lag of −0.07 seconds **(Fig. 3b)** and during *Alert* was −0.02 ± 0.14 at 0.1 seconds **(Fig. 3c)**, indicating very little time difference between cortical gamma-band and corresponding pupil diameter fluctuations during awake behaviors. Extrema in the cross-correlation in *NREM* were: −0.26 ± 0.06 at −0.1 seconds; *REM:* 0.05 ± 0.04 at 1.53 seconds; *Asleep:* −0.32 ± 0.05 at −0.43 seconds; and *All:* −0.23 ± 0.11 at −0.2 seconds.

**Fig. 3.**
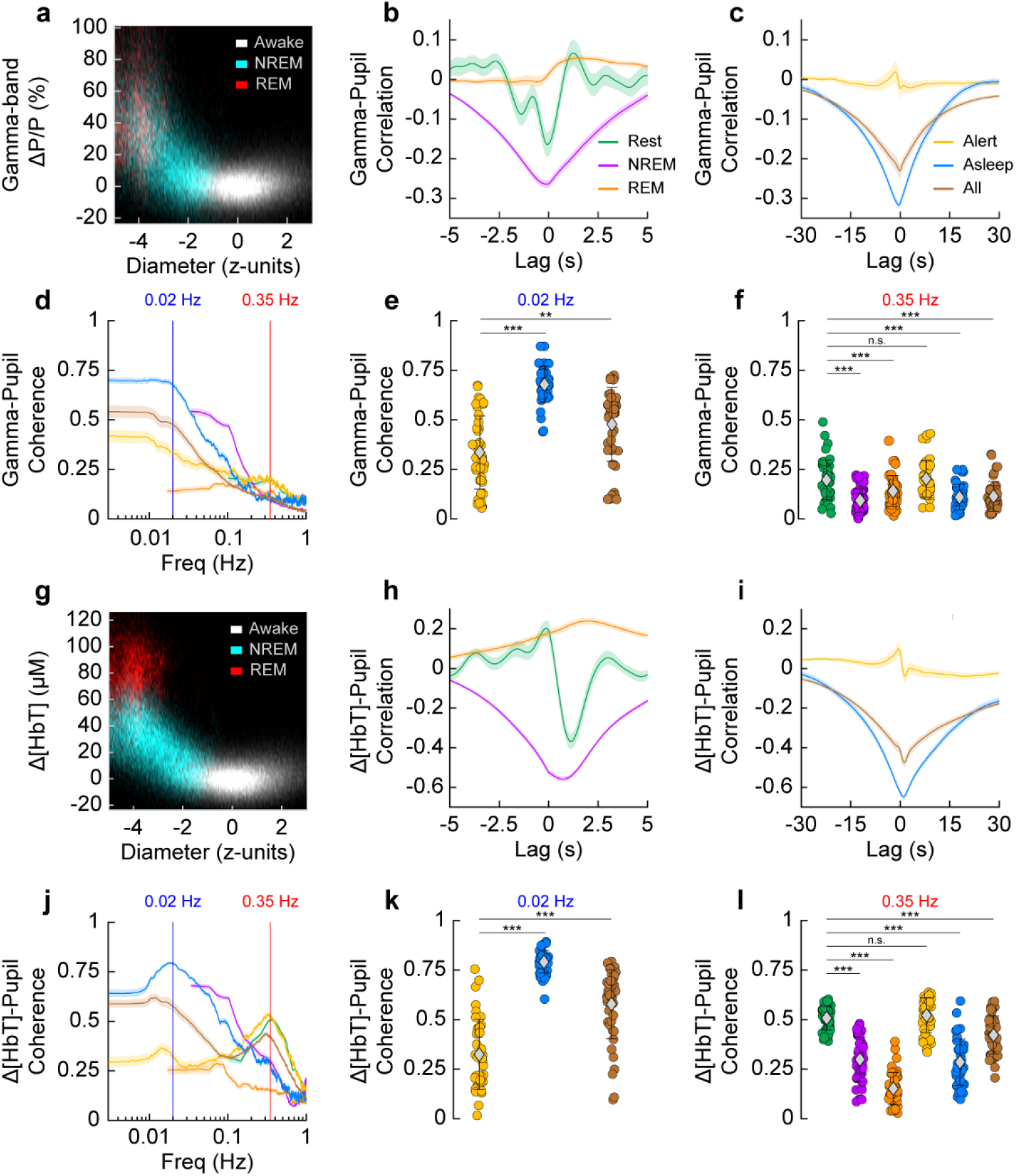
Pupil diameter shows an arousal state dependent negative correlation with blood volume and gamma-band power. **(a-f)** Relationship between gamma-band power and pupil diameter. **(a)** 2-D histogram showing the relationship between pupil diameter and gamma-band power during the three arousal state classes. **(b)** Cross-correlation between gamma-band power and pupil diameter for short duration arousal states. **(c)** Cross-correlation for longer duration arousal states. **(d)** Coherence between gamma-band power and pupil diameter. **(e)** Coherence at 0.02 Hz between gamma-band power and pupil diameter. **(f)** Coherence at 0.35 Hz. **(g-l)** Relationship between Δ[HbT] and pupil diameter **(g)** 2-D histogram showing the relationship between pupil diameter and blood volume during the three arousal state classes. **(h)** Cross-correlation between pupil diameter and blood volume for short duration arousal states. **(i)** Cross-correlation for long duration arousal states. **(j)** Coherence between pupil diameter and blood volume. **(k)** Coherence at 0.02 Hz. **(l)** Coherence at 0.35 Hz. Shading **(b, c, h, i)** denotes standard error of the mean. Error bars **(d, e, f, j, k, l)** denote standard deviation. Statistics comparisons shown are between *Rest/Alert* and other arousal states using a GLME mode with Bonferroni correction for multiple comparisons (3 in **e**, **k** *α < 0.017, ** < 0.003, *** < 0.0003 or 15 in **f**, **l**, * < 0.003, **α < 0.00067, *** < 0.000067, n.s. – not significant). See **Table 3**, **Table 4**, **Table 5**, and **Table 6** for additional statistical comparisons between each arousal state in **(e, f, k, l)**

**Fig. 4.**
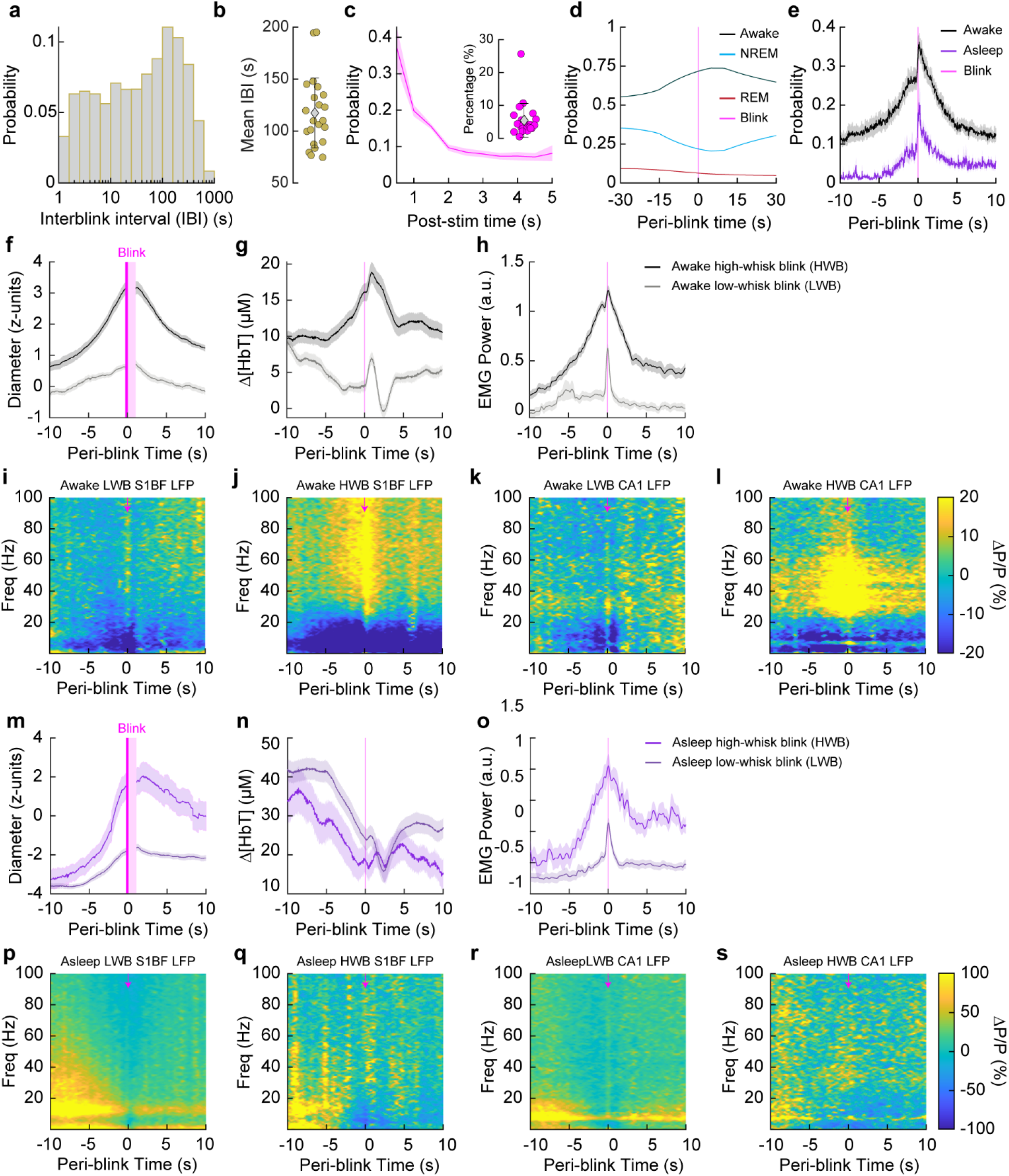
Blink-triggered neural and blood volume responses show arousal state-dependent changes. **(a)** Histogram of inter-blink interval for all blinks. **(b)** Mean inter-blink interval from each animal. **(c)** Histogram of post-stim probability of a blink for whisker stimulations that elicited a blink. (Inset) Mean probability from each animal of blinking after a whisker stimulus. **(d)** Peri-blink arousal states. **(e)** Peri-blink probability of whisking during ‘Awake’ or ‘Asleep’ periods. **(f-l)** Blink-triggered averages for blinks that occurred during an ‘Awake’ period, separated into either high whisk blinks (HWB) or low whisk blinks (LWB). Periods where the pupil is partially obscured by the eyelid have been censored (pink) **(f)** Pupil diameter during ‘Awake’ blinks **(g)** Hemodynamics during ‘Awake’ blinks **(h)** Electromyography during ‘Awake’ blinks. **(i)** Cortical LFP during ‘Awake’ low-whisk blinks (LWB). **(j)** Cortical LFP during ‘Awake’ high whisk blinks (HWB). **(k)** Hippocampal LFP during ‘Awake’ LWBs. **(l)** Hippocampal LFP during ‘Awake’ HWBs. **(m-s)** Blink-triggered averages for blinks that occurred during an ‘Asleep’ period, separated into either HWB or LWB events. **(m)** Pupil diameter during ‘Asleep’ blinks **(n)** Hemodynamics during ‘Asleep’ blinks **(o)** Electromyography during ‘Asleep’ blinks. **(p)** Cortical LFP during ‘Asleep’ blinks with low amounts of whisking. **(q)** Cortical LFP during ‘Asleep’ blinks with high amounts of whisking. **(r)** Hippocampal LFP during ‘Asleep’ LWBs. **(s)** Hippocampal LFP during ‘Asleep’ HWBs. Scatter plot error bars **(b, c inset)** are standard deviation. Shading **(c main, e, f, g, h, m, n, o)** denotes standard error of the mean.

**Fig. 5.**
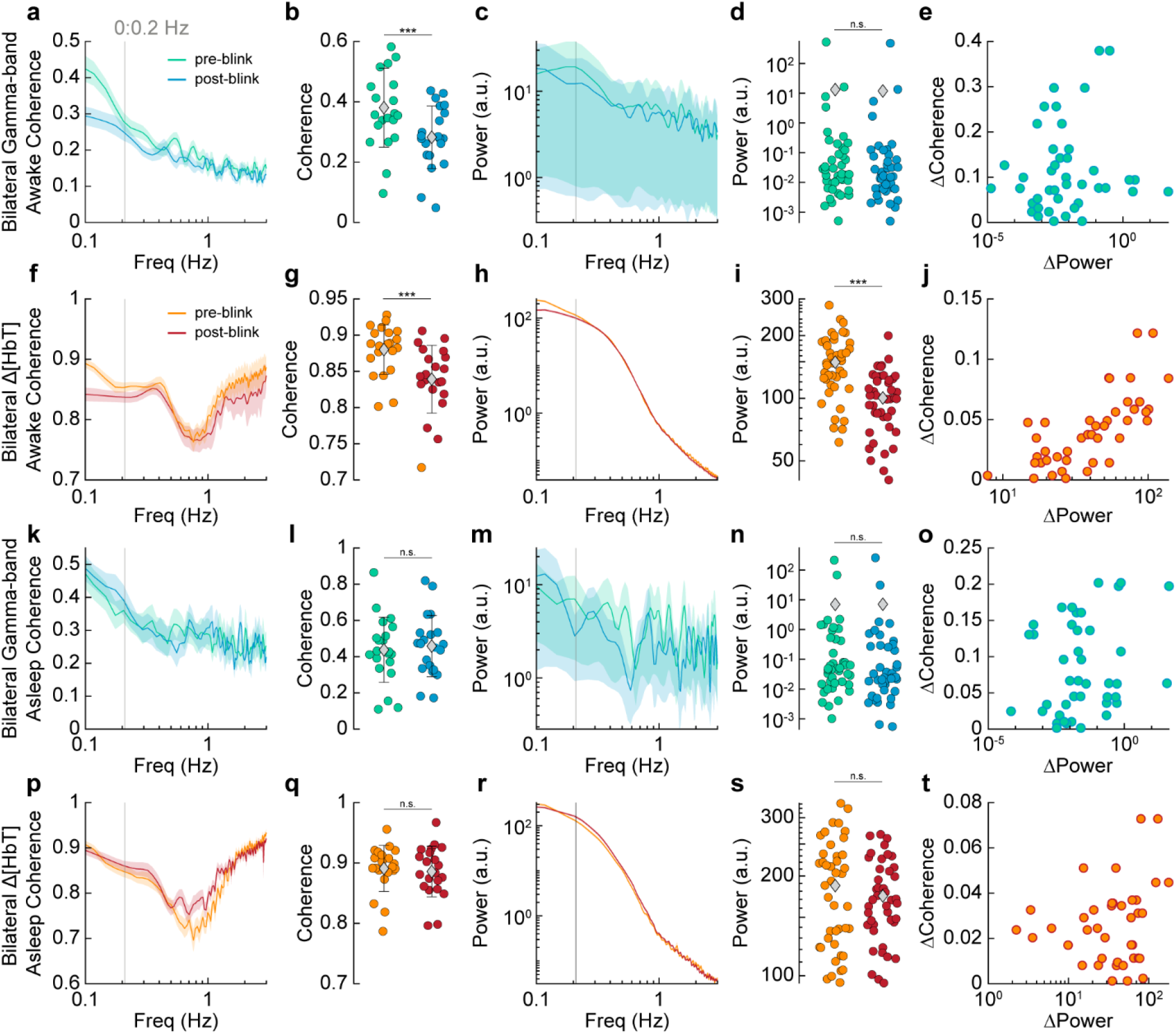
Effects of blinking on neural and bilateral hemodynamic signals. **(a)** Coherence between left and right gamma-band power before and after an awake blink. **(b)** Mean coherence from 0 to 0.2 Hz before and after an awake blink. **(c)** Power spectrum of gamma-band power envelope before and after an awake blink. **(d)** Mean power between 0 to 0.2 Hz before and after an awake blink. **(e)** Change in pre- vs. post-blink gamma-band power vs. change in pre- vs. post-blink gamma-band coherence for awake blinks. **(f)** Coherence between left and right Δ[HbT] before and after and awake blink. **(g)** Mean coherence between left and right Δ[HbT] signals between 0 to 0.2 Hz before and after an awake blink. **(h)** Δ[HbT] power before and after an awake blink. **(i)** Mean Power between 0 to 0.2 Hz before and after an awake blink. **(j)** Change in pre- vs. post-blink power vs. change in pre- vs. post-blink coherence for awake blinks. **(k)** Coherence between bilateral gamma-band signals before and after an asleep blink. **(l)** Mean coherence between bilateral gamma-band signals between 0 to 0.2 Hz before and after an asleep blink. **(m)** Power spectrum of the gamma-band power envelope before and after an asleep blink **(n)** Mean power in the gamma band envelope between 0 to 0.2 Hz before and after an asleep blink. **(o)** Change in pre- vs. post-blink power vs. change in pre- vs. post-blink coherence for asleep blinks. **(p-t)** Δ[HbT] before and after an asleep blink **(p)** Coherence between bilateral hemodynamic signals before and after a blink during sleep. **(q)** Mean coherence between 0 to 0.2 Hz before and after an asleep blink. **(r)** Δ[HbT] power before and after an asleep blink. **(s)** Mean power between 0 to 0.2 Hz before and after an asleep blink. **(t)** Change in pre- vs. post-blink power vs. change in pre- vs. post-blink coherence for asleep blinks. Error bars **(b, d, g, i, l, n, q, s)** are standard deviation. Shading in **(a, c, f, h, k, m, p, r)** denote standard error of the mean. Paired t-test *α < 0.05, **α < 0.01, ***α < 0.001, n.s. – not significant.

We next evaluated the coherence between gamma-band power and pupil diameter during different arousal states (using GLME models) and at two different frequencies. The specific frequencies (0.02, 0.35 Hz) were chosen for statistical comparison as they were the approximate peaks in pupil-gamma power coherence during the *Asleep* and *Alert* behavioral states. Using the Bonferroni correction for multiple comparisons (3 at 0.02 Hz, 15 at 0.35 Hz) put the adjusted significance thresholds (α) at 0.017 and 0.003 respectively. The pupil-gamma power coherence during *Alert* periods at 0.02 Hz was 0.34 ± 0.18 **(Fig. 3d, e, Table 3)**, and was significantly elevated during periods of *Asleep* at 0.68 ± 0.09 (*p* < 2.4×10^-18^) and during *All* data at 0.48 ± 0.19 (*p* < 2.9×10^-5^). Pupil-gamma power coherence at 0.35 Hz **(Fig. 3d, f, Table 4)** during periods of awake *Rest* and *Alert* were similar, at 0.2 ± 0.1 and 0.2 ± 0.09, respectively (*p* < 0.58), similar to what is seen between the pupil diameter and cortical neuron membrane potential (McGinley *et al.*, 2015). The coherence of all other arousal states at 0.35 Hz were significantly lower than those of the non-sleep states; *NREM:* 0.09 ± 0.06 (*p* < 2.9×10^-13^); *REM:* 0.14 ± 0.08 (*p* < 3.7×10^-5^); *Asleep:* 0.11 ± 0.07 (*p* < 9.8×10^-9^); *All*: 0.11 ± 0.08 (*p* < 5.3×10^-9^).

**Table 3.**
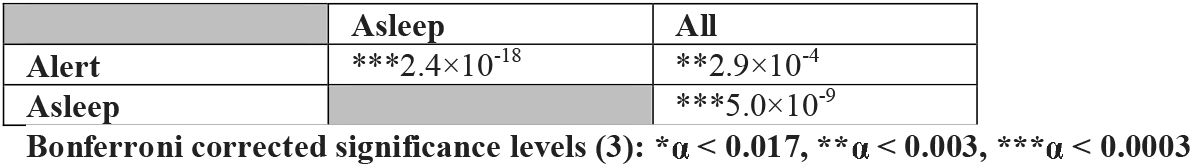
Statistical comparisons of pupil-gamma power coherence at 0.02 Hz.

|  | Asleep | All |
| --- | --- | --- |
| Alert | ***$2.4 \times 10^{-18}$ | **$2.9 \times 10^{-4}$ |
| Asleep | | ***$5.0 \times 10^{-9}$ |

Bonferroni corrected significance levels (3): \* $\alpha < 0.017$ , \*\* $\alpha < 0.003$ , \*\*\* $\alpha < 0.0003$

**Table 4.** Statistical comparisons of pupil-gamma power coherence at 0.35 Hz.

|  | NREM | REM | Alert | Asleep | All |
| --- | --- | --- | --- | --- | --- |
| Rest | ***$2.9 \times 10^{-13}$ | ***$3.7 \times 10^{-5}$ | 0.58 | ***$9.8 \times 10^{-9}$ | ***$5.3 \times 10^{-9}$ |
| NREM | | **0.0005 | ***$3.2 \times 10^{-14}$ | 0.12 | 0.10 |
| REM | | | ***$5.6 \times 10^{-6}$ | 0.07 | 0.07 |
| Alert | | | | ***$1.2 \times 10^{-9}$ | ***$6.2 \times 10^{-10}$ |
| Asleep |  |  |  |  | 0.96 |

Bonferroni corrected significance levels (15): \* $\alpha < 0.003$ , \*\* $\alpha < 0.00067$ , \*\*\* $\alpha < 0.000067$

We then looked at how the relationship between pupil diameter and cortical blood volume changed with arousal state. During the ‘Awake’ state in the absence of stimulation, fluctuations in pupil diameter and hemodynamic signals were small. When the animals were in ‘REM’ and ‘NREM’ sleep, the pupil constricted, while blood volume increased substantially (**Fig. 3g**). We then looked at the cross-correlation between Δ[HbT] and pupil diameter to determine their temporal relationship. We saw the extrema of the cross-correlation during awake *Rest* was −0.37 ± 0.25 at a lag of 1.13 seconds, such that pupil diameter changes occurred on average 1.13 seconds before hemodynamic oscillations, and pupil diameter was negatively correlated with blood volume **(Fig. 3h)**. *NREM* was more strongly anti-correlated than *Rest* at - 0.56 ± 0.08 at 0.73 seconds. *REM* had a maximum positive correlation of 0.24 ± 0.11 at 1.97 seconds. The peak of the Δ[HbT] and pupil diameter correlation in the *Alert* condition was 0.1 ± 0.21 at −0.43 seconds; this negative lag is likely due to sustained bouts of whisking **(Fig. 3i)**. *Asleep* and *All* were anti-correlated at −0.65 ± 0.08 at a lag time of 1.0 seconds, and −0.48 ± 0.15 at a lag of 1.23 seconds respectively. Generally, pupil dilations preceded vasoconstriction by about a second.

**Table 5.** Statistical comparisons of pupil-Δ[HbT] coherence at 0.02 Hz.

|  | Asleep | All |
| --- | --- | --- |
| Alert | ***$2.0 \times 10^{-29}$ | ***$9.3 \times 10^{-14}$ |
| Asleep | | ***$1.3 \times 10^{-10}$ |

Bonferroni corrected significance levels (3): \* $\alpha < 0.017$ , \*\* $\alpha < 0.003$ , \*\*\* $\alpha < 0.0003$

For nearly all cases, the coherence between hemodynamic signals and pupil diameter was substantially higher at lower frequencies. We evaluated the pupil-Δ[HbT] coherence during different arousal states at 0.02 Hz **(Fig. 3j, k, Table 5)** and 0.35 Hz **(Fig 3j, l, Table 6)**. As with the comparisons between pupil diameter and gamma-band power, these specific frequencies were chosen for statistical comparison (GLME) as they were the approximate peaks in pupil-Δ[HbT] coherence during *Asleep* behaviors and *Alert* behaviors, respectively. A Bonferroni correction for multiple comparisons (3 comparisons at 0.02 Hz, 15 at 0.35 Hz) put the adjusted significance threshold (α) at 0.017 and 0.003, respectively. At 0.02 Hz, coherence between total hemoglobin and pupil diameter during the *Alert* state was 0.32 ± 0.18 (N = 20 mice). Since we can only evaluate the 0.02 Hz component of the spectrum in the longer 15-minute duration epochs, statistical comparisons only done for longer duration behavioral conditions. Pupil-Δ[HbT] coherence during *Asleep* periods, was 0.79 ± 0.06 (*p* < 2.0×10^-29^) and *All* periods was 0.58 ± 0.17 (*p* < 9.3×10^-14^). At 0.35 Hz, *Rest* had a pupil-Δ[HbT] coherence of 0.51 ± 0.06 (N = 22 mice). The pupil-Δ[HbT] coherence during rest was not significantly different than that during the *Alert* behavior, being 0.52 ± 0.09 (*p* < 0.33). However, the coherence at 0.35 Hz was significantly lower during all sleep associated periods – *NREM:* 0.3 ± 0.11 (*p* < 7.0×10^-33^); *REM:* 0.15 ± 0.08 (*p* < 9.8×10^-66^); *Asleep:* 0.29 ± 0.12 (*p* < 2.5×10^-33^); *All*: 0.42 ± 0.1 (*p* < 3.4×10^-8^).

**Table 6.** Statistical comparisons of pupil-Δ[HbT] coherence at 0.35 Hz.

|  | NREM | REM | Alert | Asleep | All |
| --- | --- | --- | --- | --- | --- |
| **Rest** | ***7.0$\times 10^{-33}$ | ***9.8$\times 10^{-66}$ | 0.33 | ***2.5$\times 10^{-33}$ | ***3.4$\times 10^{-8}$ |
| **NREM** | | ***2.6$\times 10^{-19}$ | ***6.9$\times 10^{-35}$ | 0.59 | ***1.4$\times 10^{-14}$ |
| **REM** | | | ***1.0$\times 10^{-66}$ | ***7.7$\times 10^{-17}$ | ***7.1$\times 10^{-47}$ |
| **Alert** | | | | ***2.6$\times 10^{-35}$ | ***3.9$\times 10^{-10}$ |
| **Asleep** | | | | | ***1.9$\times 10^{-15}$ |

Bonferroni corrected significance levels (15): \* $\alpha < 0.003$ , \*\* $\alpha < 0.00067$ , \*\*\* $\alpha < 0.000067$

Low-frequency fluctuations in blood volume are highly correlated with the pupil diameter, suggesting that the same processes that modulate arousal on these timescales also cause constriction in the cerebral vasculature. There was a strong negative correlation between pupil diameter and blood volume as well as between pupil diameter and gamma-band power during periods of *Rest* and *Asleep,* with these correlations turning positive during periods of alertness and active whisking. This is in general consistent with an anticorrelation between arousal level and blood volume (Cardoso et al., 2019) in the absence of stimulation or active whisking behavior.

### Blinking is associated with increases in arousal

We next explored the relationship between blinking and arousal level. The mean inter-blink interval was 117.4 ± 33.6 seconds (**Fig. 4a, b**, N = 22 mice). We then quantified the probability of a blink being elicited by sensory stimulation of the whiskers. Except for one animal, our mice did not frequently blink after a whisker stimulation, with the mean probability of blinking within 5 seconds post-stimulus being only 5.4 ± 5.1% (**Fig. 4c**, N = 21 mice, excluded mouse at 25.6%). For blinks that did occur following stimulation, blinks typically occurred within a half-second (~37%) and the probability decreased thereafter. Blinks primarily occurred during the ‘Awake’ state (> 70%), but when they did occur during periods of ‘REM’ or ‘NREM’ sleep the animal either quickly returned to sleep after a brief awakening or did not wake up at all **(Fig. 4d)**. Blinking events also occurred during periods of maintained REM sleep with no awakening. The likelihood of volitional whisking occurring simultaneously with blinking behavior was high **(Fig. 4e)**, with volitional whisking occurring during 40% of ‘Awake’ blink events. Whisking was less frequent during blinks that occurred either during or immediately following periods classified as ‘Asleep’ (with the majority being in ‘NREM’).

As blinks commonly occurred in clusters, for all blink-related analysis we only looked at the first blink occurring in a string of blink events that occurred within less than 1 second of each other. All blinks that occurred within ± 5 seconds of whisker stimulation were also excluded to prevent the stimulus from interfering with the blink-triggered comparison. To help separate the effect of the blink from the effects’ of volitional whisking behavior coinciding with it, we split the data set into low-whisk blinks (LWB) and high-whisk blinks (HWB), further split by whether they occurred when the mouse was ‘Awake’ or ‘Asleep’ (‘REM’ and ‘NREM’ combined). LWB were defined as events where the mouse was whisking for less than 1/3 of a second in the 2 seconds surrounding each blink, whereas HWB were defined as whisking for greater than 1 second within ± 2 seconds of the blink. We evaluated blink-triggered averages for changes in diameter (z-units), Δ[HbT], EMG power, cortical LFP, and hippocampal LFP across all blinks from all mice that met the criteria. During the ‘Awake’ state, LWB caused an increase of around 0.7 z-units in pupil diameter compared to resting baseline, while HWB caused an increase in excess of 3 z-units **(Fig. 4f)**. Δ[HbT] during LWB increased around 4 μM from the period prior to the blink, whereas HWB increased around 10 μM, probably due to the increased whisking in the time preceding the blink **(Fig. 4g)**. EMG power increases ~0.6 orders of magnitude during LWB compared to 1.2 orders of magnitude during HWB, with power increases starting about 0.5 seconds before the blink occurs. There were negligible increases in gamma-band power in both the somatosensory cortex **(Fig. 4i)** and the hippocampus **(Fig. 4k)** during LWB, with the ~20% increases in power seen during high whisking likely being due to the whisking (Winder *et al.*, 2017) and not the blink **(Fig. 4j, l)**.

Blink-associated neural and vascular changes were similar during blinks that occurred during periods of ‘Asleep’, with the caveat that the baseline pupil diameter was lower. Pupil diameter during both high- and low-whisk asleep blinks was small prior to the blink (< −3 z-units), with the diameter slowly increasing preceding the blink as the animal began to wake up. For periods of low whisking, the pupil remained constricted as the animals presumably fell back asleep or remained asleep **(Fig. 4m)**. Changes in hemodynamics followed a similar trend, with there being a reduction in total hemoglobin in the time preceding the blink as the animals are waking up **(Fig. 4n)**. EMG power changes were similar to the ‘Awake’ blink, but with a lower baseline power **(Fig. 4o)**. We noted no appreciable differences among the blink-associated power spectra during blinks that occurred around sleeping periods **(Fig. 4p, q, r, s)**, however the drop in cortical delta power around the blink is evident as the animals wake up from sleep (which was predominantly NREM). Altogether, blinking was associated with increases in arousal, but to varying degrees marked by differing amounts of accompanying whisking. When blinks occurred during the ‘Awake’ state, there was only a small increase in blood volume if there was minimal whisking. However, when blinks occurred during the ‘Asleep’ state, large decreases in blood volume followed regardless of whisking amount.

### Awake blinking causes a resetting of neural and vascular dynamics

We next asked how blinking is correlated with the neural and vascular synchrony between bilateral regions of somatosensory cortex. Processing of new or startling information is accompanied by blinking, suggesting it is associated with some sort of mental ‘resetting’ (Siegle et al., 2008). As before, we separated blinks by arousal state (‘Awake’, ‘Asleep’) and looked at the changes in power and coherence during periods preceding (−15 seconds to −5 seconds) and following (5 seconds to 15 seconds) each blink. We excluded the time ± 5 seconds adjacent to each blink to capture the pre- and post-effects of the blink on synchrony and not the blink itself, as the blink elicits brief changes in neural activity and blood volume, and this response will drive coherence across all frequencies temporally close to the blink that reflect the evoked activity, not necessarily a state change in the brain. For both bilateral cortical gamma-band power signals and bilateral changes in Δ[HbT], we evaluated the power/coherence changes from 0 to 0.2 Hz as this corresponded to the frequency domain of the largest difference between pre- and post-blink. One animal was excluded from the analysis in **Fig. 4** due to being an outlier in its blink rate (N = 21). Coherence between bilateral gamma-band power signals during the ‘Awake’ state dropped following a blink from 0.38 ± 0.13 (pre-blink) to 0.28 ± 0.10 (post-blink) (*p* < 0.001, paired t-test) **(Fig. 5a, b)**. Power in these signals **(Fig. 5c)** did not change in the period following a blink, 13.2 ± 81.0 a.u. (pre-blink) versus 11.9 ± 73.7 a.u. (post-blink) (*p* < 0.25, paired t-test**) (Fig. 5d)**. There was no clear relationship between changes in gamma-band power pre- vs. post-blink vs. changes in bilateral gamma-band coherence pre- vs. post-blink **(Fig. 5e)**. ‘Awake’ bilateral Δ[HbT] coherence was 0.88 ± 0.03 (pre-blink) dropping to 0.84 ± 0.05 (post-blink) (*p* < 6.4×10^-6^) **(Fig. 5f, g)**. Unlike the gamma-band power, there was a significant drop in cortical hemodynamic power following a blink: 149.1 ± 49.1 a.u. (pre-blink) vs. 100.4 ± 33.3 a.u. (post-blink) (*p* < 8.2×10^-13^, paired t-test) **(Fig. 5h, i)**, and these appear to be related **(Fig. 5j)**.

For blinks that occurred during the ‘Asleep’ state, neither the gamma-band power nor total hemoglobin showed any significant changes between pre- and post-blink power or coherence. The mean coherence from 0-0.2 Hz between bilateral gamma-band signals during ‘Asleep’ blinks was 0.44 ± 0.18 (pre-blink) versus 0.46 ± 0.17 (post-blink) (*p* < 0.38, paired t-test) **(Fig. 5k, l)** with power changes of 7.0 ± 34.4 (pre-blink) versus 7.1 ± 39.9 (post-blink) (*p* < 0.98, paired t-test) **(Fig. 5m, n)** with no relationship between power and coherence changes **(Fig. 5o)**. Bilateral hemodynamic coherence during ‘Asleep’ blinks was 0.89 ± 0.04 (pre-blink) versus 0.89 ± 0.04 (post-blink) (*p* < 0.45, paired t-test) **(Fig. 5p, q)** at a power of 187.1 ± 67.1 (pre-blink) versus 174.1 ± 47.3 (post-blink) (*p* < 0.17, paired t-test) **(Fig. 5r, s)** with no relationship between power and coherence changes **(Fig. 5t)**. These results indicate that blinking during the ‘Awake’ state was correlated with a ‘resetting’ of neural and vascular signals, but this did not happen with blinks during ‘Asleep’.

### Pupil size, position, and motion change with arousal state

When looking at the probability of being in a given arousal state as a function of pupil diameter **(Fig. 6a),** it is apparent that the probability of wakefulness decreases dramatically as the pupil decreases in size with the 50% probability (that is, equal probability of being awake or asleep) being around −2 z-units from the resting baseline. However, the pupil size in our data set was similar across *REM* and *NREM* sleep states. We next asked how other eye metrics could help differentiate further between REM and NREM sleep, as previous work has found systematic changes in eye position of rodents during REM (Sanchez-Lopez and Escudero, 2011). When the animals fall asleep, the pupil begins to drift as the muscles around the eye relax. There are also ‘rapid-eye-movements’ characteristically seen during REM sleep. We tracked the centroid of the pupil **(Fig. 6b)** and changes in its position measured relative to the resting location. When looking at the eye velocity in each arousal state **(Fig. 6c)**, we see that the pupil moves significantly more during ‘REM’ sleep (0.66 ± 0.41 mm/sec) than in either the ‘Awake’ (0.19 ± 0.05 mm/sec, *p* < 1.8×10^-5^) or ‘NREM’ states (0.22 ± 0.15 mm/sec, *p* < 1.5×10^-5^). Despite a large change in diameter, there was minimal change in pupil centroid location during transitions between the ‘Awake’ and ‘NREM’ states **(Fig. 6d, e)**. However, the centroid of the pupil moves between 0.1 and 0.2 mm in the nasal and ventral directions when the animal transitions into the ‘REM’ state **(Fig. 6f)** and quickly returns to the baseline location upon awakening **(Fig. 6g)**.

**Fig. 6.**
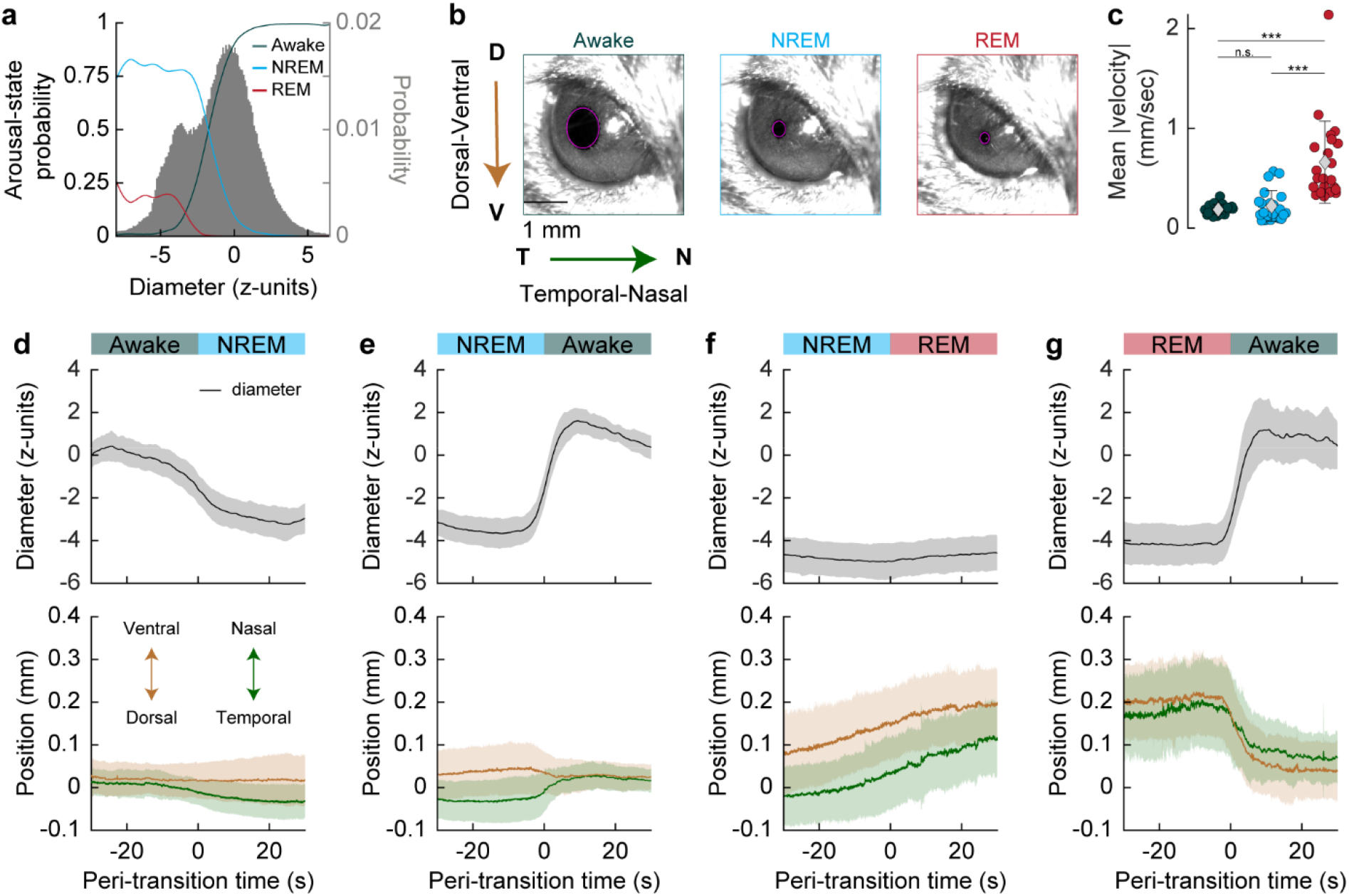
Pupil size, position, and motion change with arousal state. **(a)** Probability of arousal state classification as a function of pupil diameter **(b)** Diagram demonstrating the drift of the pupil as the animal falls to sleep. The position of the centroid is measured relative to the resting location. **(c)** Mean absolute velocity of the eye in each arousal state. **(d-g)** Pupil diameter (top) and pupil position (bottom) as the animal transitions from one arousal state to another. During REM sleep, the eye rotated towards the nose (nasally) and ventrally. Error bars and shading denote standard deviations. Bonferroni corrected (3) paired t-test *α < 0.017, ** < 0.003, ***α < 0.0003, n.s. – not significant.

### Eye metrics are an accurate predictor of arousal state

Finally, we explored using eye metrics (pupil diameter and position) alone as a predictor of arousal state, comparing their predictive power to that of more conventional sleep scoring using physiological parameters (cortical delta power, hippocampal theta power, EMG). As pupil diameter is commonly measured in behaving mouse paradigms (International Brain et al., 2021; McGinley *et al.*, 2015; Musall et al., 2019; Pisauro *et al.*, 2016; Stringer et al., 2019; Vinck *et al.*, 2015), arousal scoring using pupil diameter and eye movement could be very useful for detecting bouts of sleep when other more invasive assays are not available. Previous work has shown that the sleep/wake state of head-fixed mice can be determined accurately on the timescale of hundreds of seconds using pupil diameter (Yuzgec et al., 2018), but here we asked if it can be done on a time scale of seconds.

Pupil diameter alone could be used to differentiate the ‘Awake’ arousal state from the ‘REM’ and ‘NREM’ states (with a threshold at approximately −2 z-units, **Fig. 6a**). However, there was much less difference in pupil diameter between the two sleep states. Therefore, we used the position and motion of the pupil’s centroid in our model classification to achieve a separation between ‘REM’ and ‘NREM’. We compared the manual scores of an exemplar 15-minute imaging session to those produced by three different bootstrapped random forest classification models **(Fig 7a)** following model training using a 70:30 training:testing split. The first model utilized only eye metrics, such as the pupil’s diameter (mm, z), position, and average velocity (‘Eye Model’) (see Materials and Methods). We used the same example data shown in **Fig. 1f** to show how these three metrics can be used to identify arousal state (**Fig. 7b, c, d**). The pupil decreases in size during ‘REM’ and ‘NREM’ sleep as compared to the awake state, but it is the changes in pupil position that separate ‘REM’ from ‘NREM’ sleep. The second model was identical to that used in (Turner et al., 2020) and used seven physiological parameters (i.e., ‘Physiological Model’, see Materials and Methods). The third model was a combination of the two (‘Combined Model’), utilizing both eye metrics and the physiological measurements together. With respective to identifying sleep, the Eye Model (N = 22 mice) had a total cumulative accuracy of 90.4% across all testing data **(Fig. 7e)** with comparable type-I and type-II classification errors. The Physiological Model (cumulative accuracy of 93.0%) **(Fig. 7f)** and combined model (cumulative accuracy of 93.9%) **(Fig. 7g)** performed better than the eye model, having access to the “gold-standard” of electrophysiology and electromyography information classically used for detecting sleep. The out-of-bag error during model training was higher for the Eye Model (0.10 ± 0.02) than the Physiological Model (0.08 ± 0.02, *p* < 7.0×10^-6^), as well for as the Combined Model (0.07 ± 0.02, *p* < 5.2×10^-10^) **(Fig. 7h)**. However, the eye metrics did significantly improve the Combined Model over the original Physiological Model (*p* < 1.0×10^-6^). Our findings indicate that monitoring the eye, including pupil diameter and its changes in position, can be used as a non-invasive method to determine if head fixed mice are sleeping on a timescale of seconds.

**Fig. 7.**
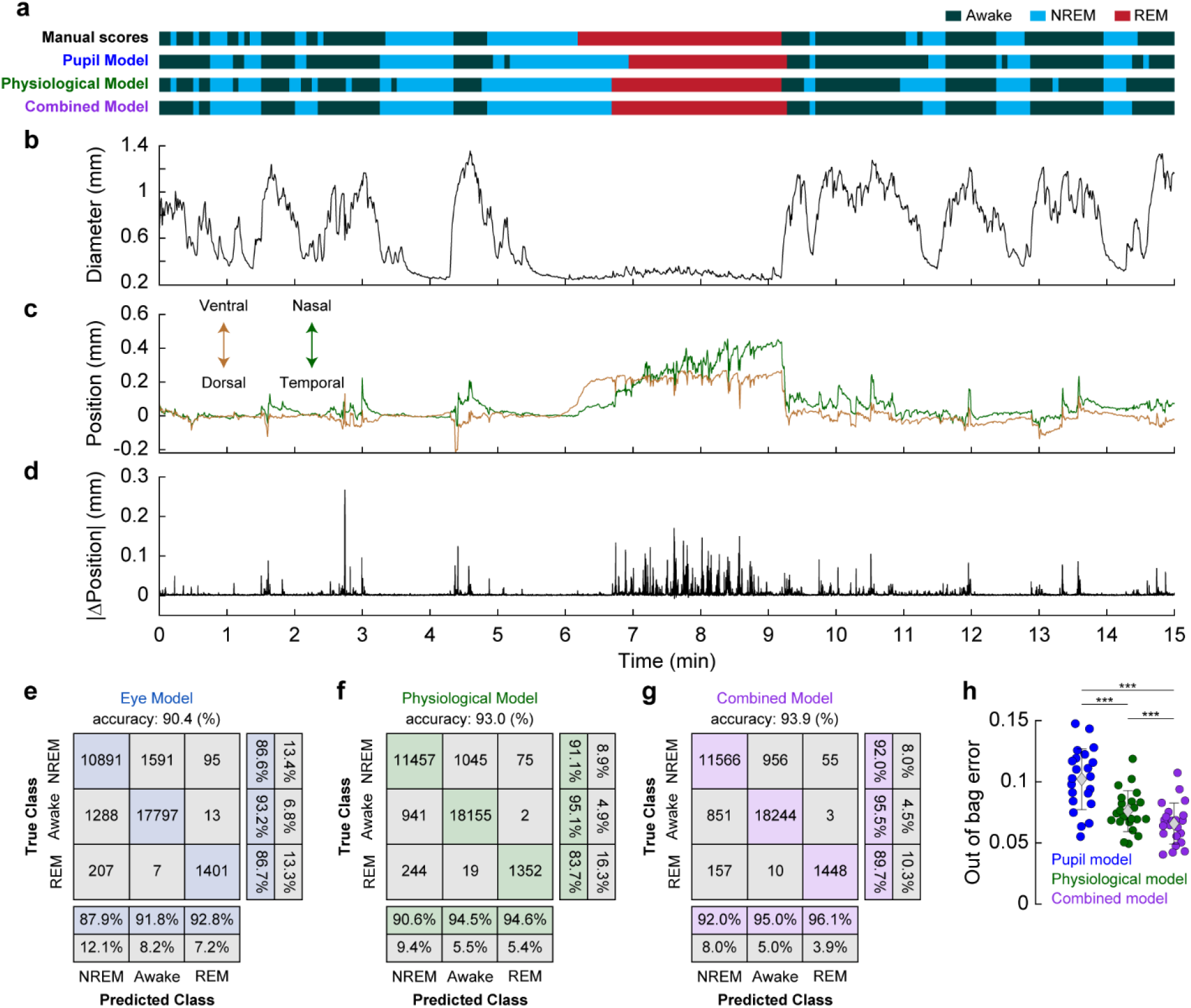
Eye metrics alone are an accurate predictor of arousal state. **(a)** Arousal state predictions from 3 arousal state classification models (Eye, Physiological, Combined) and manual scores for sample data. Data is the same as presented in Figure 1. **(b)** Example trace of pupil diameter over time with respect to arousal state. For clarity, blinks are not shown. **(c)** Position of the pupil changes when the animal goes into ‘REM’ sleep, moving nasally and ventrally. **(d)** Absolute change in the pupil’s frame-by-frame position indicates a high degree of movement during ‘REM’ sleep. **(e)** Confusion matrix of a bagged random forest (70:30% training:testing) using only eye metrics such as pupil diameter, position, and motion (‘Eye Model’) **(f)** Confusion matrix of a bagged random forest (70:30% training:testing) using measurements of cortical and hippocampal LFP, electromyography, heart rate, and whisker motion (‘Physiological Model’). **(g)** Confusion matrix of a model using both the eye and physiological data (‘Combined Model’) for arousal scoring. **(h)** Out-of-bag error estimation during model training for each model. Error bars in **(h)** denote standard deviation. Bonferroni corrected (3) paired t-test *α < 0.017, **j < 0.003, ***α < 0.0003, n.s. – not significant.

## Discussion

In this study, we explored the relationship between pupil diameter and blinking with ongoing neural activity and hemodynamic signals in the somatosensory cortex of mice during different arousal and behavioral states. Pupil diameter was consistently smaller during sleep states and larger during the awake state, allowing the pupil to be used as a predictor of arousal state (Yuzgec et al., 2018). As a practical matter, a threshold of approximately 2 z-units below the resting pupil diameter can function as an indicator of sleep. Blinking was correlated with changes in arousal levels, as well as with neural and vascular dynamics. Mice were more likely to be in the awake state after blinking than before, and bilateral synchronization in the gamma-band envelope and accompanying hemodynamics both decreased after blinks when the mouse was awake. The difference in neural responses to blinks in the awake versus asleep conditions could be due to differences in sensory processing in these respective states.

Because our mice slept frequently, when averaged over the entire data set the pupil diameter was strongly anti-correlated with both gamma-band power and blood volume due to the large vasodilation occurring during NREM sleep (Turner et al., 2020). When restricted to just data in the awake state (whose neural and hemodynamic signals are dominated by body movements and fidgeting behavior (Drew et al., 2020; Drew et al., 2019; Huo et al., 2014; Musall et al., 2019; Salkoff et al., 2020; Stringer et al., 2019; Tran et al., 2018; Winder et al., 2017), the pupil diameter was positively correlated with gamma-band power and blood volume **(Fig. 8)**. The coherence between blood volume and pupil diameter was stronger at lower frequencies and was highest (> 0.75) at 0.02 Hz (corresponding to a ~50 second period) in the sleeping mouse. Interestingly, activity in the LC and the concentration of noradrenaline during NREM sleep fluctuates on a similar timescale (Kjaerby et al., 2022; Osorio-Forero et al., 2021). As noradrenaline is vasoconstrictory (Bekar et al., 2012; Goadsby et al., 1985; Raichle et al., 1975), the fluctuations in noradrenaline during NREM may contribute to the oscillations in blood volume (Fultz et al., 2019; Turner et al., 2020) that simulations and experiments have suggested are important for clearing waste from the brain (Kedarasetti et al., 2020a; 2022; Kedarasetti et al., 2020b; van Veluw et al., 2020; Xie et al., 2013). The fact that there is vasodilation in somatosensory cortex during periods of body motion and arousal when noradrenaline levels are highest can be accounted for by the action of local vasodilatory signals from neurons during behavior (Echagarruga et al., 2020; Winder et al., 2017; Zhang et al., 2019). While our LFP measurements were conducted in the somatosensory cortex, we would expect similar dynamics in other sensory areas of the cortex, such as visual areas, which are innervated by the same LC neurons (Kim et al., 2016). The vascular dynamics may differ in the frontal cortex and other areas, which receive input from a different subset of LC neurons (Kim et al., 2016).

**Fig. 8.**
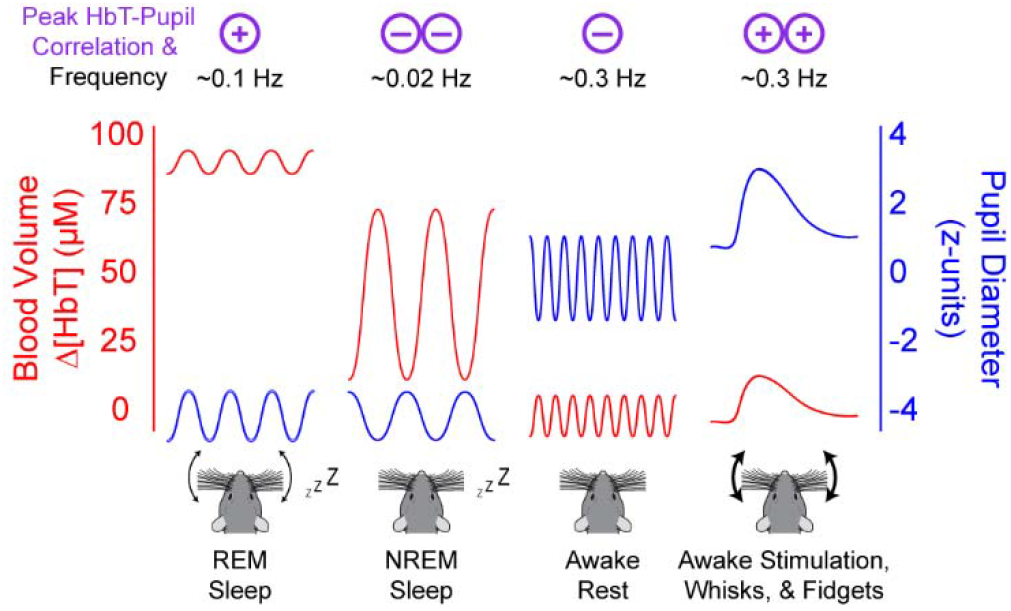
Pupil diameter is anti-correlated with cortical hemodynamics during Rest and NREM sleep. Summary schematic demonstrating the observed correlation between pupil diameter and cortical hemodynamics during each arousal state. Correlations between pupil diameter and blood volume are positive during REM sleep and when the mouse is alert, moving, or stimulated. Correlations between pupil diameter and blood volume are negative during rest and NREM sleep. While the oscillation amplitude and baseline offset of the pupil diameter/blood volume reflect what is observed in our data, for clarity the temporal frequencies of oscillations are not to scale.

Our findings that the pupil diameter was strongly anticorrelated with blood volume should be taken in the context of the literature relating arousal and pupil diameter to LC activity, and the vasoconstrictory impact noradrenergic inputs of LC on the cerebral vasculature. The noradrenaline levels in the brain correspond with LC neuron cell body activity (Berridge and Abercrombie, 1999; Feng et al., 2019; Poe et al., 2020). Sensory stimuli drive increases in LC neural activity, noradrenergic tone, and pupil dilation (Gilzenrat et al., 2010; Gray et al., 2021; Joshi et al., 2016; Murphy et al., 2014; Schwarz and Luo, 2015; Yang et al., 2021) though the correlations between pupil diameter and spiking of individual LC neurons is low (Megemont et al., 2022). Activation of noradrenaline-releasing neurons in the LC drive awakening from sleep (Carter et al., 2010; Hayat et al., 2020; Kjaerby et al., 2022), and this awakening from sleep is followed by profound cortical vasoconstriction (Turner et al., 2020) and brain-wide hemodynamic changes (Fultz *et al.*, 2019; Liu et al., 2018; Liu et al., 2015; Zhang et al., 2022a). Electrical stimulation of the LC and subsequent increases in noradrenaline levels (Bekar et al., 2012) cause vasoconstriction (Goadsby et al., 1985; Raichle et al., 1975) (but see (Toussay et al., 2013) that observe that stimulation of the LC can increase cortical perfusion in the anesthetized rat). We saw that the blood volume-pupil diameter correlation was positive in the *Alert* state, which seems to contradict the vasoconstrictory nature of noradrenaline. The *Alert* state contains many fidgeting and whisking bouts, which can be nearly continuous body movements. While cortical noradrenaline levels are known to rise during locomotion (Paukert et al., 2014; Polack et al., 2013) and movement (Feng et al., 2019), the activity of vasodilatory neuronal nitric oxide synthase (nNOS) neurons (Echagarruga et al., 2020) and vasodilatory extracellular potassium are both elevated during locomotion (Longden et al., 2017; Rasmussen et al., 2019), and these two vasodilators are likely strong enough to drive vasodilation even the face of elevated noradrenaline. Furthermore, the magnitude of noradrenaline increases during movement are much smaller than the decreases during sleep. During sleep, microdialysis measured cortical noradrenergic levels fall > 50% (Bellesi et al., 2016), though recent studies using fluorescent biosensors have shown there are large fluctuations over the timescales of minutes in noradrenaline levels in frontal areas during NREM (Kjaerby et al., 2022). The arousal-induced noradrenergic levels increases during voluntary body movements and fidgeting are of a substantially smaller magnitude than the decreases seen during sleep (Feng et al., 2019), which would explain why vasodilation dominates in these behaviors. The positive correlations between pupil diameter and blood volume during REM might be due to fluctuations in cholinergic drive, which is high during REM sleep (Jing et al., 2020; Jones, 2020) and is also linked to pupil dilations (Larsen and Waters, 2018). Our observations of state-dependent correlation between blood volume and pupil diameter (**Fig. 8**) is consistent with noradrenergic modulation in the cortex playing a state-dependent role in neurovascular coupling in concert with local neural activity (Hamel, 2006; Kleinfeld et al., 2011).

Monitoring the pupil (Privitera *et al.*, 2020) can provide a second-by-second insight into the sleep/wake state of mice. These periods of sleep can be an issue not only in behavioral tasks but can also be a confound in studies looking at spontaneous neural and hemodynamic activity. In human studies, sleep episodes occur frequently during resting-state imaging (Tagliazucchi and Laufs, 2014), drastically affecting functional connectivity measurements and other measures of network dynamics. In mice, bilateral neural and hemodynamic correlations are much higher during sleep (Turner et al., 2020), supporting the need for arousal-state monitoring in both animal and human studies as detecting and monitoring changes in arousal is essential for accurate interpretations of any resting-state study (Drew *et al.*, 2020; Liu *et al.*, 2018; Tagliazucchi and Laufs, 2014). Fortunately, monitoring pupil diameter and eye motion can provide a simple, non-invasive way of detecting sleep and monitoring arousal state transitions in rodents.

**Video 1.** Video showing pupil diameter variations across several arousal states. Detected pupil area is in purple.

## Acknowledgements

We thank Nikki Crowley for comments and feedback on the manuscript. This work was supported by NIH grants R01NS078168 and R01NS079737 to P.J.D.

## Author Contributions

K.L.T. and P.J.D. designed the experiments. K.LT performed experiments. K.L.T., K.W.G., and P.J.D. analyzed the data. K.L.T. and P.J.D. wrote the paper.

